## Supplemental Material for "A proximity complementation assay to identify small molecules that enhance the traffic of ABCA4 misfolding variants"

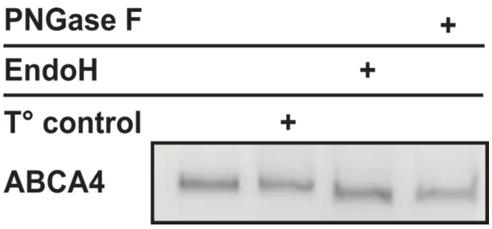


**Figure S1. Glycosylation status of ABCA4 in mouse retina.** 6 µg of protein lysate from mouse retina were treated with EndoH, PNGase-F, or buffer and temperature only protocol-control (T°). A slight increase in protein mobility was observed following both PNGase-F and EndoH. N= 2 mice retinas.

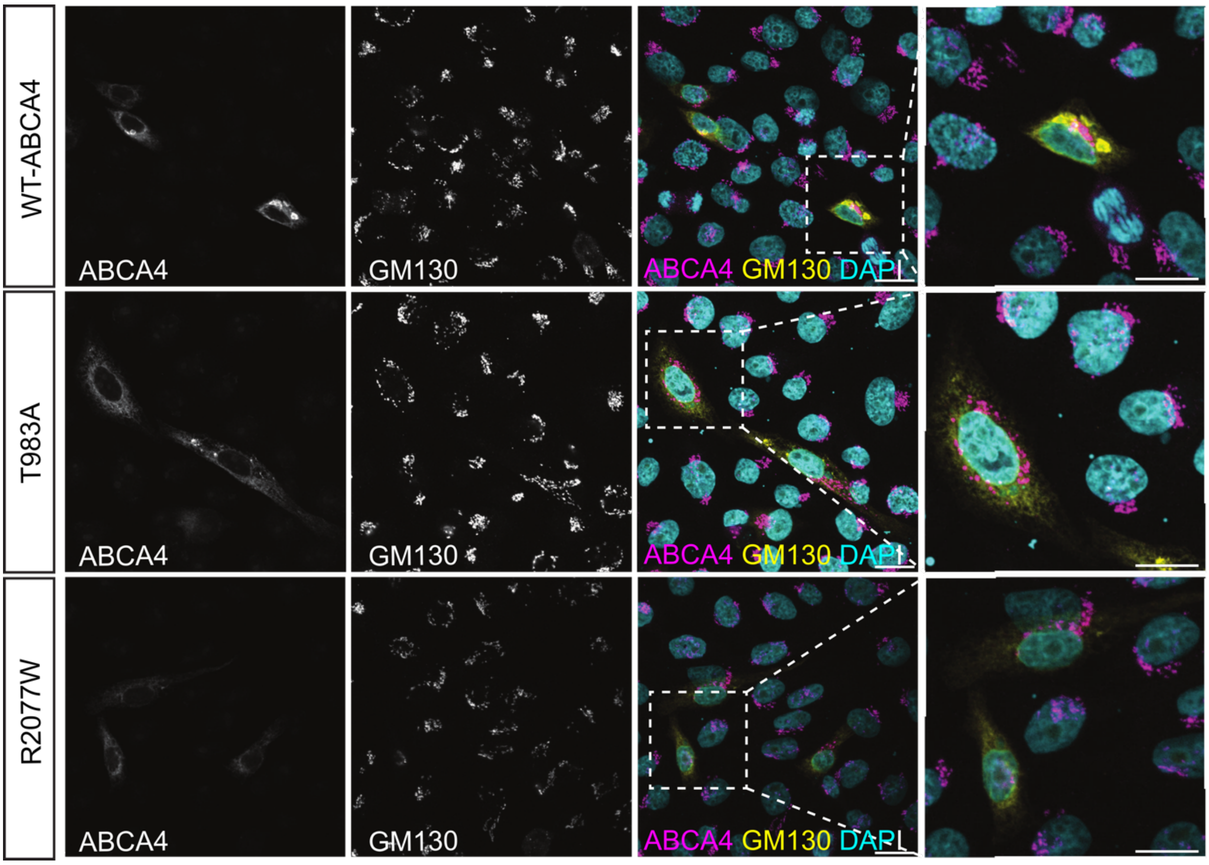


**Figure S2. ABCA4 and Golgi localization in cultured cells.** CHO cells were transfected with plasmid expressing WT ABCA4 protein and the indicated variants. 48 hours post-transfection cells were fixed in 4% PFA and permeabilised with Triton X-100 and double labelled with the ABCA4-Abbexa (yellow) and GM130 (magenta). ABCA4 was not detected overlapping with the Golgi. Scale bars = 10 µm

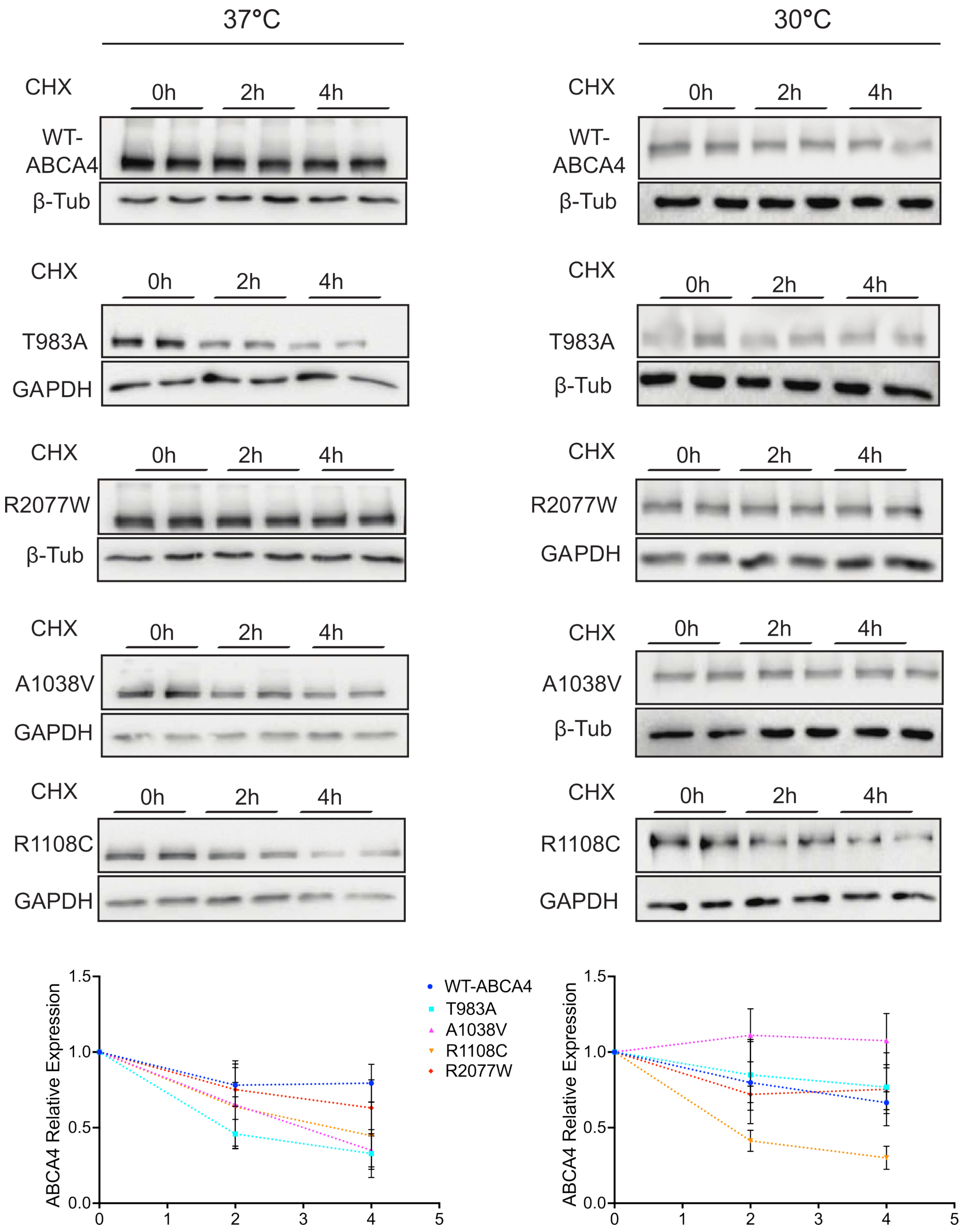


**Figure S3. The effect of the temperature on ABCA4 protein degradation.** HEK293T cells were transfected with plasmids expressing WT ABCA4 and variants and incubated at 37ºC or 30ºC. 48h post-transfection cells were treated with 50 µM of (CHX) for 0, 2 and 4 h and western blotted (10 µg of protein lysate). **A**) Quantification of protein levels by ImageJ. Data were normalised to GAPDH/β-Tubulin. Mean of fold change + SD. n = 3 independent experiments.

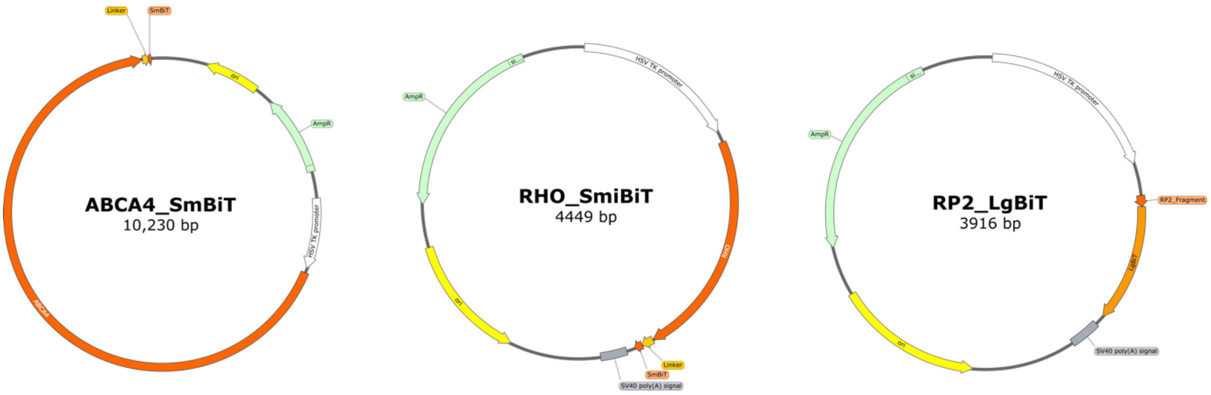


**Figure S4. Plasmid maps.** The RP2-LBiT, RHO-SmBiT and ABCA4-SmBiT plasmid maps. The P23H and the ABCA4 missense variants are not shown since they all are single nucleotide variants in the coding sequence and the vector backbones are the same.

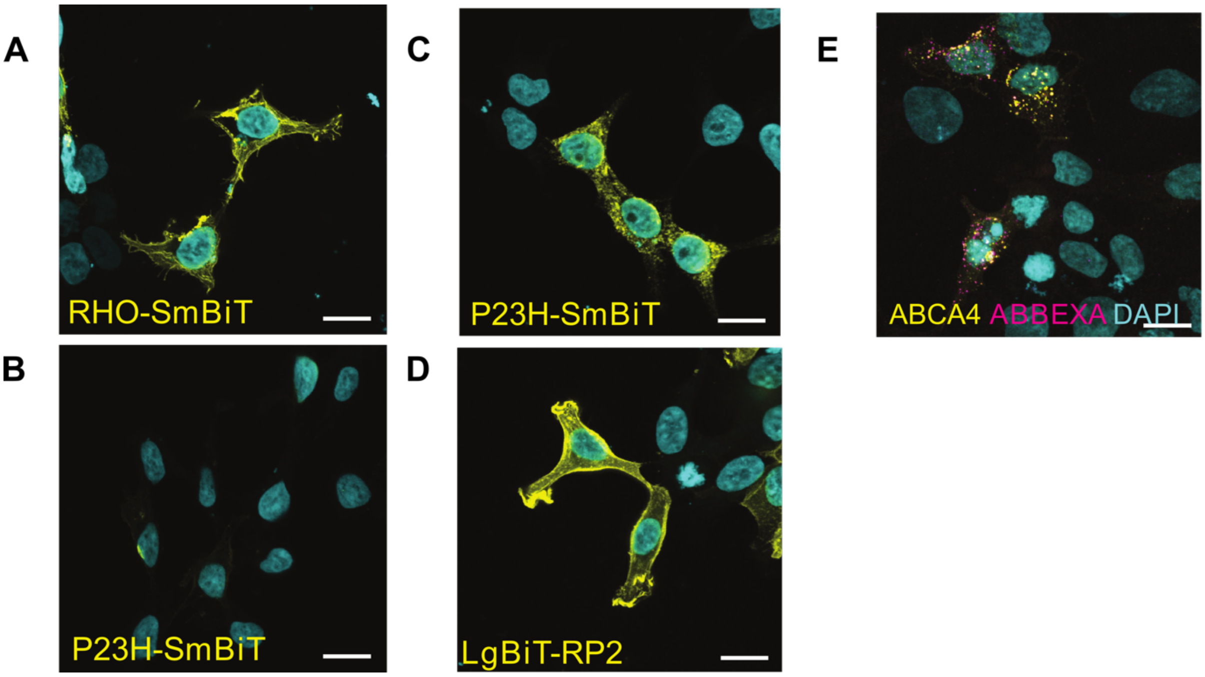


**Figure S5. Localization of nanoBiT fusion proteins.** HEK293T cells were transfected with plasmids expressing the different protein constructs. 24 hours post-transfection cells were fixed in 4% PFA and permeabilised with Triton X-100. A non-permeabilizing protocol was used when indicated **A)** WT RHO fused with SmBiT localises to the plasma membrane. The staining was performed with antibody 4D2 without a permeabilization step, as the 4D2 epitope is located on the extracellular N-terminus the protein. **B)** P23H-SmBiT is not detected with 4D2 using a no-permeabilization protocol. **C)** P23H fused to SmBiT staining is observed in a perinuclear and reticular pattern, consistent with the ER, following a detergent permeabilization step to reveal 4D2 immunoreactivity. **D)** The RP2-LgBiT localised to the plasma membrane, when stained with anti-LgBiT antibody in permeabilized cells. **E)** Intracellular WT-ABCA4-SmBiT was detected with the ABCA4 3F4 ABCAM antibody and WT-ABCA4-SmBiT localising to the plasma membrane was detected using an ABBEXA antibody 48h post-transfection analysis Scale bars = 10 µm.

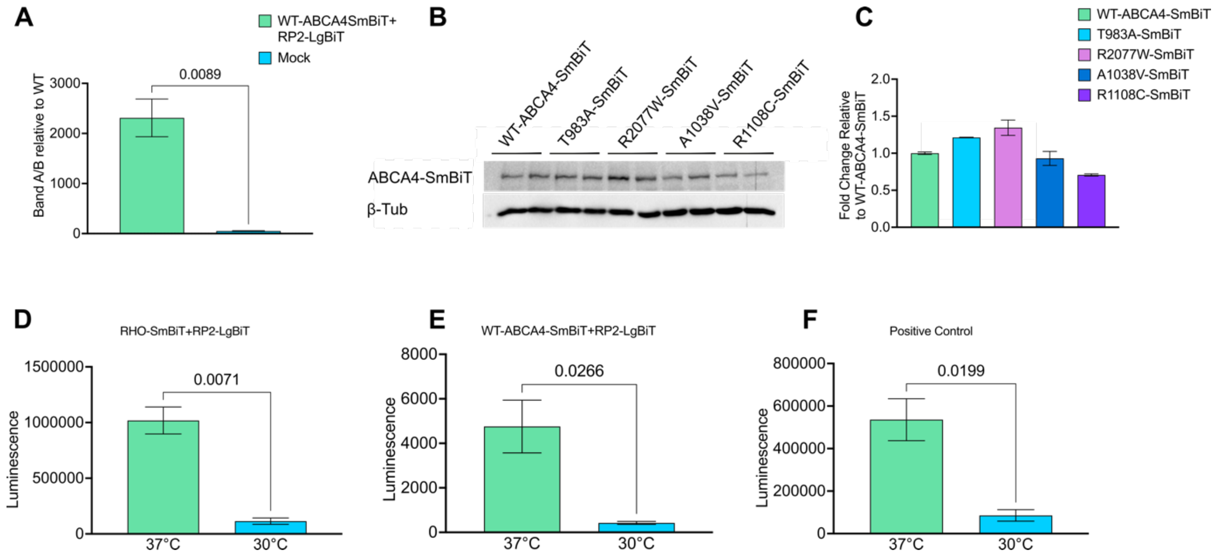


**Figure S6. Validation of split NanoBit complementation assay A**) 48h post-transfection live HEK293T cells were analysed using a luminometer. WT-ABCA4-SmBiT + RP2-SmBiT shows a stronger luminescence with compared to the non-transfected cells (Mock). Error bars are SD. n = 3. Two-tailed Student's t-test. **B,C**) 48h post-transfection HEK293T cells were collected, and the protein lysate was analysed by western blot and quantification was obtained with ImageJ. No marked decrease in the steady state level of the ABCA4 proteins was detected. n = 2. **D-F**) Luminescence produced at 37ºC was compared to the signal produced at 30ºC using the RHO-SmBiT+RP2-LgBiT and the ABCA4-SmBiT+RP2-LgBiT plasmids in HEK293T. A significant drop in luminescence signal was observed at 30ºC in HEK293T cells in both conditions. **F**) Luminescence signal at 37ºC vs 30 ºC was also studied using the positive control plasmids provided in the NanoLuc Binary Technology system by Promega. A significant drop in luminescence signal was observed at 30ºC. Error bars are SD. Two-tailed Student's t-test.

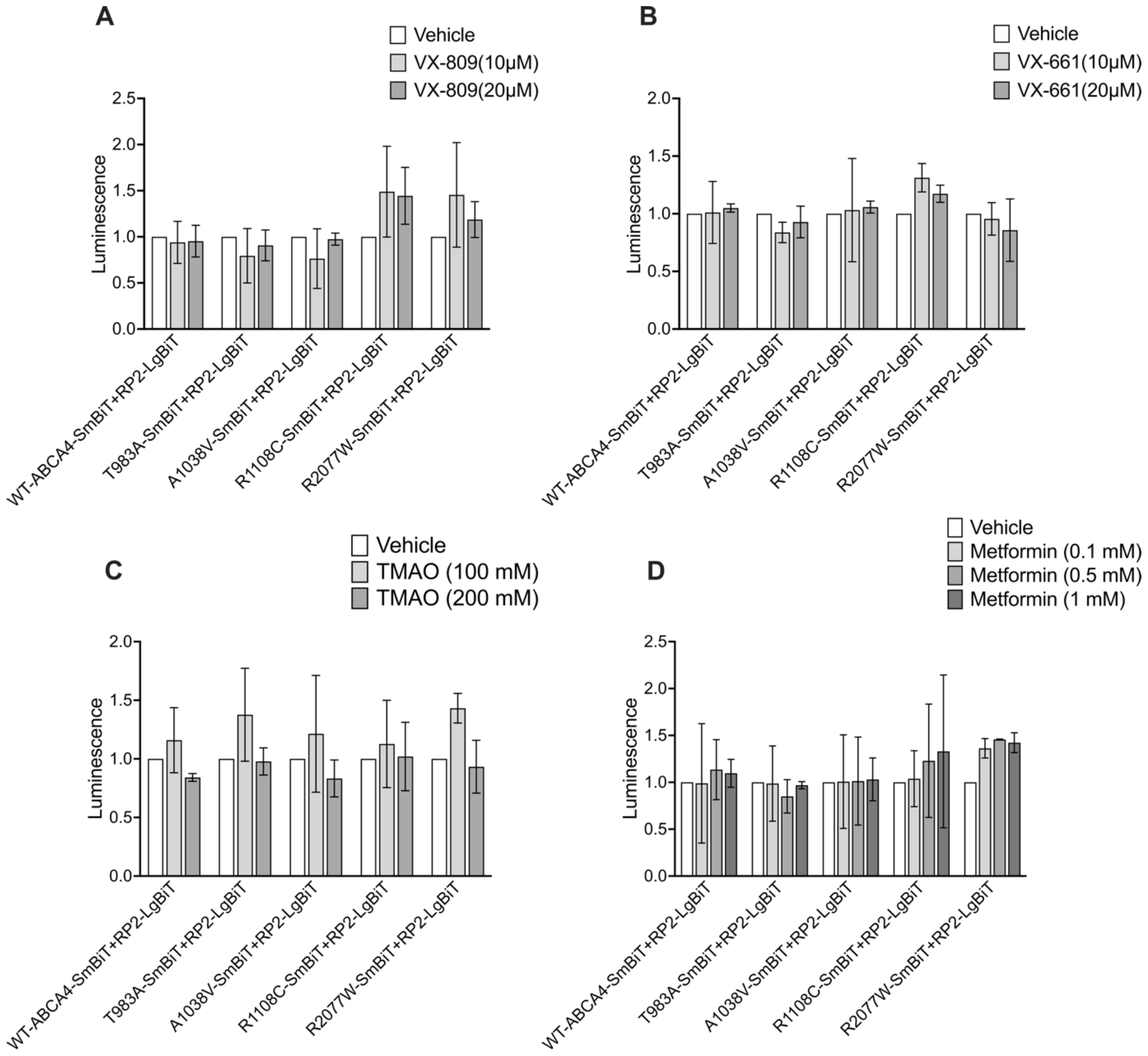


**Figure S7. Small compounds are screened using the complementation assay. A** and **B**) Luminescence signal in HEK293T was analysed after 48h from transient transfection for the indicated NanoBiT fusion combinations. 24h post transfection cells were treated with 10 µM or 20 µM of VX-809 (**A**) and VX-661 (**B**) for a total of 24h treatment. Mean fold change in luminescence relative to the vehicle + SD. n=2. **C**) 48h post transfection cells were treated with different concentrations of TMAO (100 and 200 mM) for a total of 24h treatment. Mean fold change in luminescence relative to the vehicle + SD. n=3. **D**) Luminescence signal in HEK293T was analysed after 48h from transient transfection for the indicated NanoBiT fusion combinations. 24h post transfection cells were treated with different concentration of metformin (0.1, 0.5 and 1 mM) for a total of 24h treatment. Mean fold change in luminescence relative to the vehicle + SD. n=2
